# Identification and characterization of novel bat coronaviruses in Spain

**DOI:** 10.1101/2025.01.07.631674

**Authors:** Clàudia Soriano-Tordera, Jaime Buigues, Adrià Viñals, Elena Muscolino, Raquel Martínez-Recio, Juana Díez, Juan S Monrós, José M. Cuevas, Jérémy Dufloo, Rafael Sanjuán

**Affiliations:** Institute for Integrative Systems Biology (I^2^SysBio), Universitat de València and Consejo Superior de Investigaciones Científicas, València, Spain; Institut Cavanilles de Biodiversitat i Biologia Evolutiva, Universitat de València, València, Spain; Molecular Virology group, Department of Medicine and Life Sciences, Universitat Pompeu Fabra, Barcelona, Spain; Department of Genetics, Universitat de València, València, Spain

## Abstract

The zoonotic transmission of bat coronaviruses poses a threat to human health. However, the diversity of bat-borne coronaviruses remains poorly characterized in many geographical areas. Here, we recovered six complete coronavirus genomes by performing a metagenomic analysis of fecal samples from hundreds of individual bats captured in Spain, a country with high bat species diversity. Three of these genomes corresponded to potentially novel coronavirus species belonging to the alphacoronavirus genus. Phylogenetic analyses revealed that some of these viruses are closely related to coronaviruses previously described in bats from other countries, suggesting the existence of a shared viral reservoir worldwide. Using viral pseudotypes, we investigated the receptor usage of the identified viruses and found that one of them can use human and bat ACE2, highlighting its zoonotic potential. However, the receptor usage of the other viruses remains unknown. This study broadens our understanding of coronavirus diversity and identifies research priorities for the prevention of zoonotic viral outbreaks.

**Author summary:** Bats carry many viruses, some of which can cross the species barrier and infect humans, a process known as zoonosis. In particular, bat-borne coronaviruses pose a significant threat to human health. To improve our pandemic preparedness, it is essential to characterize the diversity and zoonotic potential of bat coronaviruses. However, such research efforts have historically suffered from a strong geographical bias. For example, despite the rich bat diversity in Spain, few studies have searched for coronaviruses in Iberian bats. Here, we used viral metagenomics to test for the presence of coronaviruses in more than 200 bat samples collected across Spain. We detected six complete coronaviruses, three of which were proposed to be new viral species. We characterized their relationship to previously identified viruses and their ability to use known coronavirus receptors to enter cells, demonstrating that one virus could use human ACE2 as a receptor. Our results highlight the diversity of bat-borne coronaviruses in Spain, their zoonotic potential, and the need to better characterize coronavirus diversity worldwide.

## Introduction

Bats are taxonomically diverse mammals that represent 20% of all mammal species (1,2). Bats have been identified as natural reservoirs for numerous zoonotic viruses, including the Nipah and Hendra paramyxoviruses (3), the hemorrhagic Ebola filovirus (4), or the severe acute respiratory syndrome (SARS) and Middle East respiratory syndrome (MERS) coronaviruses (5–7). Their frequent association with viral emergence might simply reflect the diversity and abundance of bat species (8). Alternatively, it has been suggested that bats exhibit increased propensity to carry and transmit pathogens because of their distinctive metabolism associated with flight, limited immunoinflammatory responses, tendency to aggregate into populous colonies, and adaptation to peri-urban habitats (1,2,9). Given their role as viral reservoirs, there has been a concerted research effort to explore viral diversity and identify potential zoonotic viruses in bats (10–13). This has been facilitated by the development of metagenomic next-generation sequencing (mNGS) that allows exploring the unknown virosphere and enables the identification of many new viral species, including novel bat-borne viruses (14,15).

Numerous studies have suggested that bats are frequent carriers of coronaviruses (16). Coronaviruses are enveloped, single-stranded, positive-sense RNA viruses with genomes ranging from 16 to 31 kb. The family *Coronaviridae* is divided into four genera: *Alphacoronavirus* (AlphaCoV) and *Betacoronavirus* (BetaCoV), which are associated with infections in mammals, and *Gammacoronavirus* and *Deltacoronavirus* that primarily infect birds but also occasionally mammals (17,18). Seven coronaviruses have already jumped from an animal reservoir to humans (SARS-CoV, SARS-CoV-2, MERS-CoV, 229E, NL63, HKU1 and OC43). Assessing the zoonotic potential of coronaviruses identified in wildlife is therefore critical, but not an easy task. Indeed, many deposited sequences are only partial, the full virus is rarely isolated, and working with unknown pathogens requires stringent biosafety measures. One way to overcome these limitations is to use surrogate experimental systems that focus on specific steps of the viral life cycle, such as viral entry, which is mediated by the coronavirus spike glycoprotein and plays a critical role in determining viral host range and cellular tropism (19). Viral pseudotypes, in which the spike glycoprotein is incorporated into a viral vector, can be used to safely and faithfully characterize the receptor usage of a new virus and its ability to enter human cells (13,20,21).

Given the threat they pose to human health, identifying and characterizing bat-borne coronaviruses is a priority worldwide. However, there is a strong geographical bias in the metagenomic identification of bat coronaviruses. As of November 2024, 60.4% of the coronavirus sequences deposited in the Bat-associated virus database DBatVir (22) originate from Asia, while only 6.5% have been detected in European bats. Moreover, only a few studies have been carried out to search for coronaviruses in Iberian bats and none of them provided a direct *in vitro* assessment of the zoonotic potential of the identified viruses (23,24), despite the Iberian Peninsula being home to a wide diversity of bat species. Here, we used mNGS to detect coronaviruses in a large number of fecal samples from different regions of Spain. Six complete coronavirus genomes were recovered, including three potential novel species.

## Results

### Coronaviruses found in bat fecal viromes

We obtained 202 fecal samples from 23 bat species in 20 collection points across Spain (**Figure 1**). Samples were grouped into 26 pools, each containing samples from different individuals of the same species (**Table S1**). After RNA extraction and library preparation, Illumina sequencing yielded between 4.5 and 48 million raw reads per pool (**Table S2**). Quality-filtered reads were assembled *de novo*, and the resulting contigs were used to identify viral sequences. A total of 8946 viral contigs and 430 high-quality complete or nearly complete viral genomes were obtained, the vast majority of which corresponded to bacteriophages. Among animal viruses, eight classified as belonging to the family *Coronaviridae* are the subject of the present study, while other viral families were already analyzed in previous articles (25–27). These eight sequences were identified in the feces of six different bat species: *Miniopterus schreibersii, Myotis capaccinii, Myotis daubentonii, Myotis escalerai, Pipistrellus kuhlii* and *Rhinolophus hipposideros*. Six were complete and two were partial (10-28% completeness) as determined by CheckV (28), and the number of reads assigned to each virus ranged from 266 (mean coverage 12.44x) to 47,005 (mean coverage 223.73; **Table S3**). The proposed names of these viruses and sequence accession numbers are presented in **Table 1**.

**Table 1.** Features of *de novo* assembled coronavirus sequences.

| Accession number | Proposed names | Abbreviated names | Bat species | Length (nt) | Genus | Subgenus |
| --- | --- | --- | --- | --- | --- | --- |
| PQ611137 | Sarbecovirus sp. isolate RhGB02 (variant Murcia2022) | RhBetaCoV-Murcia2022 | *Rhinolopus hipposideros* | 29259 | Betacoronavirus | Sarbecovirus |
| PQ611138 | Myotis capaccinii associated Yeseras alphacoronavirus | McAlphaCoV-Yeseras | *Myotis capaccinii* | 28323 | Alphacoronavirus | Pedacovirus |
| PQ611139 | Myotis escalerai associated Sima alphacoronavirus | MeAlphaCoV-Sima | *Myotis escalerai* | 24428 | Alphacoronavirus | Pedacovirus |
| PQ611140 | Miniopterus schreibersii associated Gordo alphacoronavirus | MsAlphaCoV-Gordo | *Miniopterus schreibersii* | 29264 | Alphacoronavirus | Minunacovirus |
| PQ611141 | Alphacoronavirus sp. isolate ATG20-YN-CHN-2020 (variant Murcia2022) | MsAlphaCoV-Murcia2022 | *Miniopterus schreibersii* | 28877 | Alphacoronavirus | Minunacovirus |
| PQ611142 | Alphacoronavirus Bat-CoV/P.kuhlui/Italy/206679-3/2010 (variant Valencia2022) | PkAlphaCoV-Valencia2022 | *Pipistrellus kuhlii* | 28171 | Alphacoronavirus | Nyctacovirus |
| - | Bat coronavirus isolate BtCoV/13585-58/M.dau/DK/2014 (variant Spain2022) | MdAlphaCoV-Spain2022 | *Myotis daubentonii* | 8147* | Alphacoronavirus | Pedacovirus |
| - | Bat coronavirus HKU7 (variant Spain2022) | MsAlphaCoV-Spain2022 | *Miniopterus schreibersii* | 2953* | Alphacoronavirus | Minunacovirus |

\*incomplete virus

**Figure 1.**
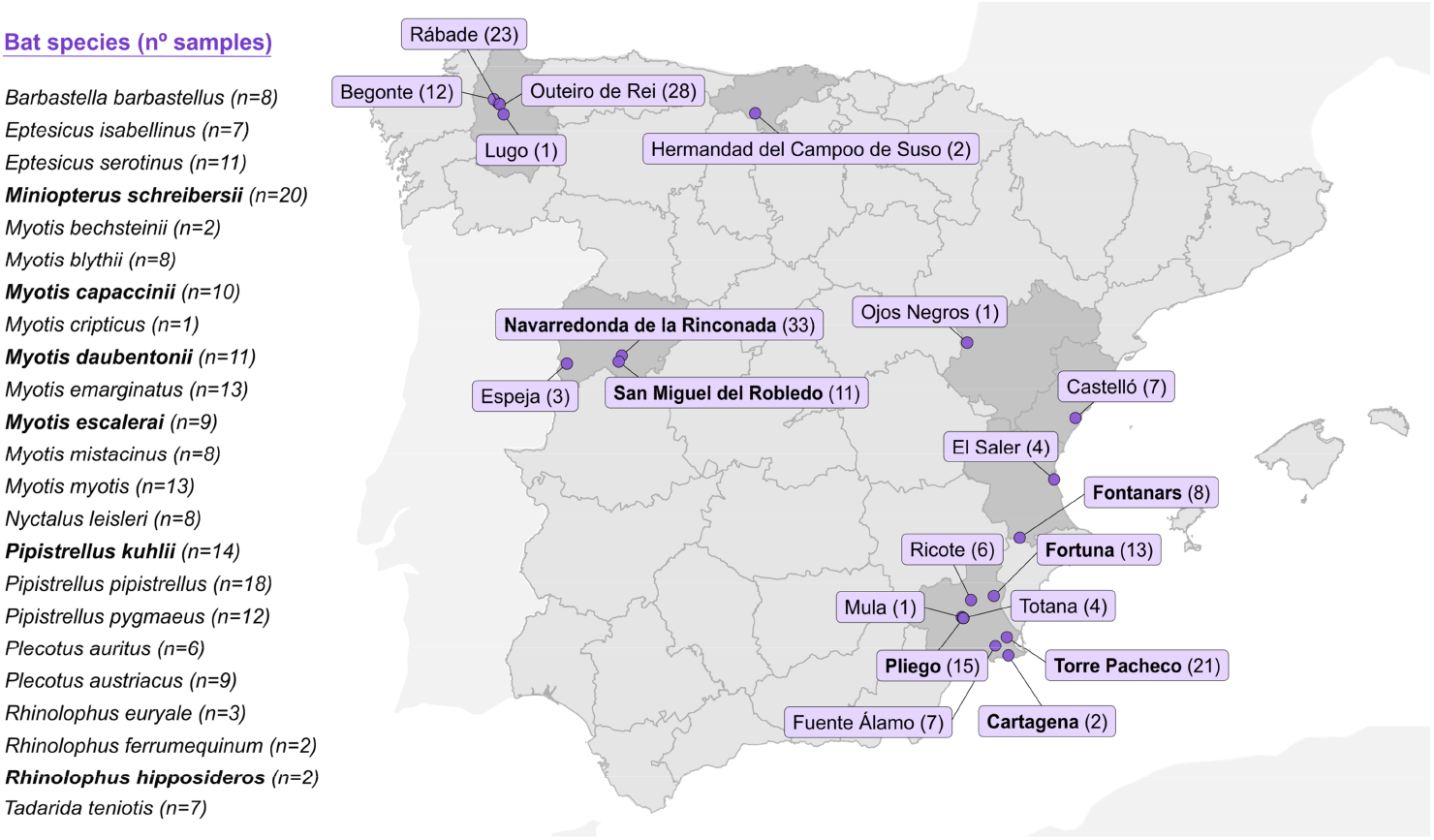
Collection of fecal samples from 23 bat species throughout Spain. The number of samples collected from each bat species and the number of individuals captured in each area are indicated in parentheses. The locations and bat species in which coronaviruses were detected are shown in bold.

### Phylogenetic and taxonomic classification of coronavirus sequences

To determine the genus and subgenus classification of the identified coronaviruses, we constructed phylogenetic trees using the RNA-dependent RNA polymerase (RdRP; **Figure 2A**), and helicase (**Figure 2B**) amino acid sequences of ICTV-approved species (18). These analyses demonstrated that seven of the eight coronaviruses were alphacoronaviruses, including three minunacoviruses (MsAlphaCoV-Gordo, MsAlphaCoV-Murcia2022 and MsAlphaCoV-Spain2022), one nyctacovirus (PkAlphaCoV-Valencia2022), and three pedacoviruses (MsAlphaCoV-Yeseras, MeAlphaCoV-Sima and MsAlphaCoV-Spain2022), whereas one was a betacoronavirus belonging to the *Sarbecovirus* subgenus (RhBetaCoV-Murcia2022).To ascertain whether these could be novel species, we determined the percent sequence identity to the closest nucleotide sequences using BLASTn. Five of the sequences showed >91% identity to previously described viruses, while three (McAlphaCoV-Yeseras, MeAlphaCoV-Sima and MsAlphaCoV-Gordo) showed <84% identity to any other known coronaviruses and thus might represent new species **(Table S3)**.

**Figure 2.**
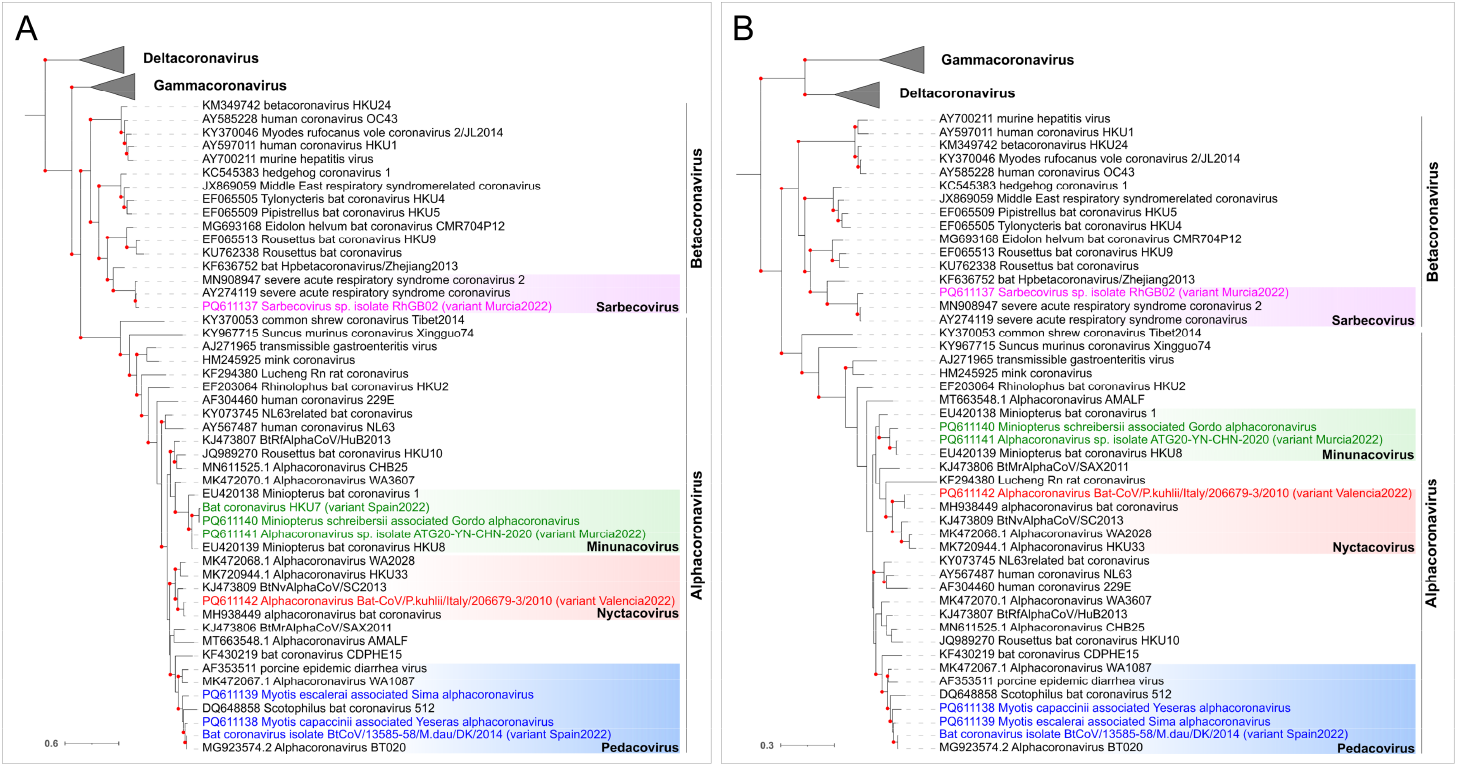
Phylogenetic positioning of the newly described viruses within the family *Coronaviridae*. Maximum-likelihood (ML) trees were built using RdRP **(A)** and helicase **(B)** amino acid sequences. Sequences from representative ICTV-approved viral species were included. The Gamma- and Deltacoronavirus taxonomic groups are collapsed by genus. Microhyla letovirus 1 was used as an outgroup to root the trees. Viruses described in this study are shown in colors. Bootstrap values higher than 80 are indicated with red circles. The scale bar indicates the evolutionary distance in amino acid substitutions per site.

### Geographical distribution and prevalence of identified coronaviruses

To determine geographical distribution and possible prevalence of the six complete coronaviruses, we used RT-PCR to detect their presence in the individual fecal samples from the pools in which they were initially identified (**Table S4**). Most corresponded to the region of Murcia. Indeed, the two pedacoviruses were identified in Pliego and Fortuna, RhBetaCoV-Murcia2022 in Cartagena and MsAlphaCoV-Murcia2022 and MsAlphaCoV-Gordo in Torre Pacheco. The only exception was PkAlphaCoV-Valencia2022, which was detected in the region of Valencia (Fontanars). In general, viruses were only detected in a single location, except for McAlphaCoV-Yeseras, which was detected in individuals from the Pliego and Fortuna regions, suggesting that this virus may be circulating in two spatially separated bat roosts. Interestingly, several coronaviruses were detected in multiple individuals from the same location, suggesting intra-roost virus circulation. A striking example was the PkAlphaCoV-Valencia2022, which was detected in 100% (8/8) of *Pipistrellus kuhlii* individuals sampled in the Fontanars collection site.

### Analysis of the closest spike sequences

To further investigate the evolutionary origin of the detected coronaviruses, spike phylogenetic trees were constructed for each subgenus using the 20 closest Blastp hits for each sequence. This analysis was done only for the six completely sequenced viruses, since the other two lacked spike reads. We found that the RhBetaCoV-Murcia2022 spike was related to another sarbecovirus, the bat SARS-like Khosta-2 (QVN46569.1) found in *Rhinolophus hipposideros* bats from Russia (29), these two sequences showing 93% identity (**Figure 3A; Table S3**). The spike of MsAlphaCoV-Gordo was related with 92% identity to another minunacovirus found in *Miniopterus schreibersii* in China (ABO88150.1) (30) (**Figure 3B; Table S3**), while the spike of the other minunacovirus, MsAlphaCoV-Murcia2022, was most closely related to a virus detected in *Rhinolophus sinicus* (UUW33738.1; 97% identity) also in China (31) (**Figure 3B; Table S3**). The spike of the nyctacovirus identified in *Pipistrellus kuhlii* (PkAlphaCoV-Valencia2022) was highly similar (up to 98% identity) to the spike of viruses found in the same bat species in Italy (YP009744890.1 and AZF86130.1) (32) (**Figure 3C; Table S3**). The spike of the pedacovirus from *Myotis capacinii* (McAlphaCoV-Yeseras) formed a clade with those of viruses found in China (WCC63178.1, WCC63185.1, WCC63171.1, WCC63164.1 and WCC62859) (33) and in Russia (ASV51733.1) (34), but shared less than 78% identity with any of these sequences (**Figure 3D; Table S3**). For the other pedacovirus spike, MeAlphaCoV-Sima, the closest spike sequence was obtained from one virus detected in *Myotis myotis* in Switzerland (USF97421.1) (35) (80% identity) (**Figure 3D; Table S3**).

**Figure 3.**
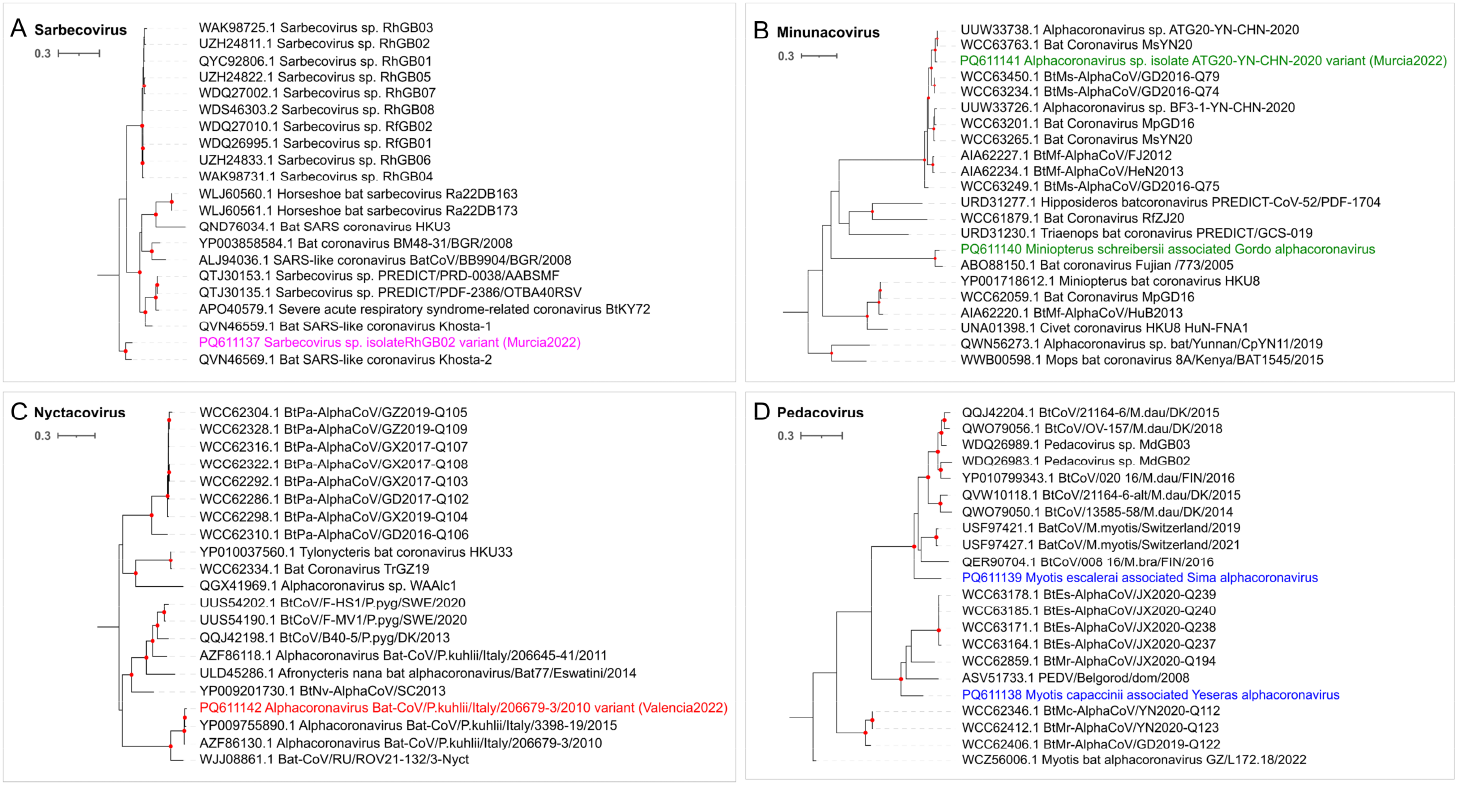
Spike sequences in the context of *Sarbecovirus (A), Minunacovirus (B), Nyctacovirus (C)* and *Pedacovirus (D)* phylogenies. ML trees were built using the 20 Blastp hits for each identified spike sequence. Viruses described in this study are shown in colors. Bootstrap values higher than 80 are indicated with red circles. The scale bar indicates the evolutionary distance in amino acid substitutions per site.

### Failure to isolate the newly identified coronaviruses

To functionally characterize these newly identified viruses, we attempted to isolate the three novel alphacoronaviruses (McAlphaCoV-Yeseras, MeAlphaCoV-Sima and MsAlphaCoV-Gordo) and the sarbecovirus (RhBetaCoV-Murcia2022) from PCR-positive fecal samples. Briefly, Huh-7 and VeroE6 cells were inoculated with positive samples in the presence or absence of trypsin, and viral load was followed by RT-qPCR. However, after four days, no significant viral replication could be measured. A second passage was attempted for the McAlphaCoV-Yeseras and the RhBetaCoV-Murcia2022 but was also unsuccessful.

### RhBetaCoV-Murcia2022 can use human and bat ACE2 to enter cells

In the absence of viral isolates, we used vesicular stomatitis virus (VSV)-based pseudotyping to explore the receptor usage of the six complete viruses identified. We successfully constructed the six viral pseudotypes, as shown by detection of spike incorporation into viral particles by Western blot (**Figure 4A**). We then tested whether these pseudotypes could enter cells overexpressing the human (h) orthologues of the known coronavirus receptors ACE2, APN, DPP4 and TMPRRS2. The expression of each receptor was also verified by Western blot (**Figure 4A**), except for hACE2 and hDPP4, whose expression was confirmed by infection with SARS-CoV-2 and MERS-CoV pseudotypes, respectively (**Figure 4B**). Interestingly, RhBetaCoV-Murcia2022 pseudotypes efficiently entered hACE2-expressing cells (**Figure 4B**), consistent with previous results obtained with the related Khosta-2 spike (36). The other five viruses could not enter cells expressing any of the four human receptors tested. However, given that interspecies variation in receptor proteins can strongly affect the ability of a virus to use them for entry (36,37), we sought to determine whether the identified spikes could use various bat ACE2, APN or TMPRSS2 orthologues. Unfortunately, none of these bat genomes have been sequenced, except for *Pipistrellus kuhlii*. To address this limitation, we managed to obtain novel mRNA sequences from bat feces samples for ACE2 from *Miniopterus schreibersii*, APN from *Myotis escalerai* and *Miniopterus schreibersii* and TMPRSS2 from *Myotis escalerai* and *Miniopterus schreibersii*. For the other gene-bat species combinations, RT-PCR from fecal samples was unsuccessful and we thus resorted on available sequences from the related *Myotis myotis, Rhinolophus sinicus* and *Rhinolophus cornutus* species. Sequences were synthesized and correctly expressed, as shown by Western blot (**Figure 4A**). Moreover, as a functional control, we verified that the ACE2 orthologues of *Myotis myotis, Miniopterus schreibersii, Rhinolophus sinicus* and *Rhinolophus cornutus* allowed SARS-CoV-2 spike-mediated entry, as previously shown (38) (**Figure 4B**). We found that, in addition to using hACE2, the RhBetaCoV-Murcia2022 pseudotype could enter cells overexpressing the ACE2 orthologues from *Myotis myotis, Miniopterus schreibersii* and *Rhinolophus sinicus* (**Figure 4B**). However, the other five viruses could not use any ACE2, APN or TMPRSS2 orthologue tested. Therefore, the receptor usage of these viruses remains unclear.

**Figure 4.**
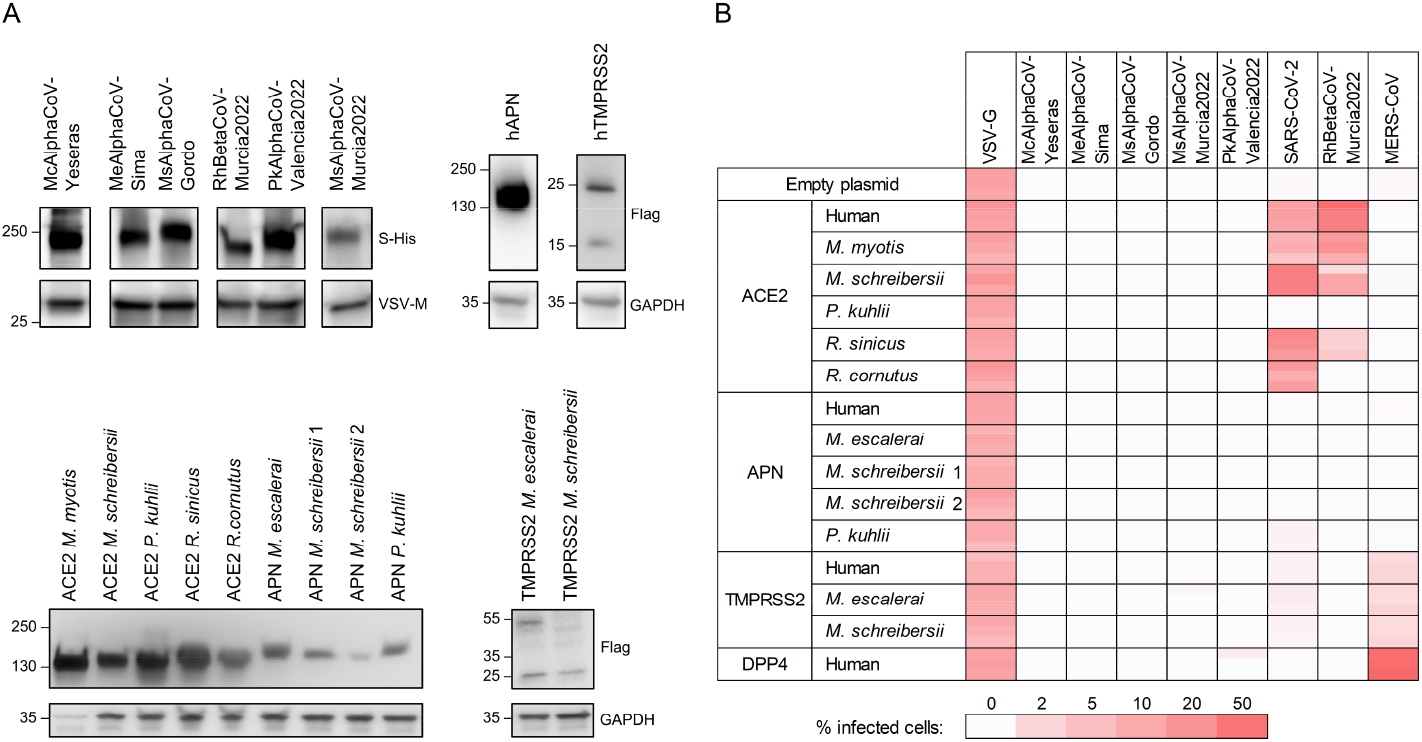
Receptor usage of the identified coronaviruses. **A**. Western blot validation of spike incorporation into VSV pseudotypes and receptor expression. The spike was detected with an anti-His-Tag antibody and VSV-M was used as a loading control, whereas receptors were detected with an anti-Flag-Tag antibody and GAPDH was used as a loading control. **B**. Heat map showing the percentage of infected cells after transfection of the indicated bat or human orthologues and inoculation with pseudotyped viruses carrying the indicated spike proteins. For each combination triplicate (n = 3) assays are shown. The two APN alleles present in *M. schreibersii* were considered.

## Discussion

We have characterized the feces virome of 23 different bat species from various regions of Spain and identified six complete and two incomplete coronavirus genomes, including three candidate novel species. Of the eight identified viruses, five exhibited close genetic similarity to previously described coronaviruses identified in the same or different bat species from distant geographic regions, particularly in Asia and other parts of Europe. For instance, variants of RhBetaCoV-Murcia2022 were detected in the same bat species (*Rhinolophus hipposideros*) in Russia (29) and in the United Kingdom, as well as MsAlphaCoV-Murcia2022 isolated in *Miniopterus schreibersii* was previously obtained from another bat species (*Rhinolophus sinicus*) in China (31). This suggests a broad geographical distribution of some bat-borne coronaviruses, potentially facilitated by bats migratory patterns and habitat overlap across the continent. Moreover, the fact that bats from different families harbour variants of the same viral species highlights the broad species tropism of some bat-borne coronaviruses. Such inter-family or inter-genus cross-species transmission of alpha- and betacoronaviruses was shown to occur frequently in Asia and America (39,40) and our data suggest that this may also be the case in Europe. Finally, the identification of three novel bat-borne coronavirus species expands our knowledge of coronavirus diversity in Europe, particularly in Spain, where studies searching for bat-borne coronaviruses have been limited (23,24).

Although metagenomic studies allow an in-depth characterization of the unknown virosphere in wildlife, experimental characterization of newly identified viruses is critical to assess their zoonotic potential. Given the presumed importance of spike-mediated entry in coronavirus cross-species transmission, we used spike-expressing viral pseudotypes to assess the receptor usage of the six complete coronaviruses identified. We showed that RhBetaCoV-Murcia2022 can use hACE2 as a receptor. Although this raises concerns about its potential zoonotic risk, this does not imply that the virus is currently capable of infecting humans. Indeed, additional incompatibilities with host proteases, immune response, or post-entry blocks in viral replication may prevent RhBetaCoV-Murcia2022 from replicating in human cells (1,16). To better examine their zoonotic potential, we tried to isolate the three new alphacoronavirus (McAlphaCoV-Yeseras, MeAlphaCoV-Sima and MsAlphaCoV-Gordo) and the RhBetaCoV-Murcia2022 betacoronavirus from PCR-positive samples, but this was unsuccessful. This could be due to low viral loads in samples, the absence of infectious virus despite the presence of viral RNA, the choice of non-susceptible cell lines for isolation, or a suboptimal dose of trypsin used, among other potential causes. Therefore, further experiments are needed to assess the ability of these viruses to replicate in human cells.

Most of the coronaviruses we identified could not use any of the tested bat orthologues of ACE2, APN or TMPRSS2. The only exception was the RhBetaCoV-Murcia2022, which could use the ACE2 orthologues of *Myotis myotis, Miniopterus schreibersii* and *Rhinolophus sinicus*, but not from *Rhinolophus cornutus*, despite a 95.3% protein identity between these last two species. Previous studies have demonstrated that minimal interspecies sequence variations among receptor proteins can deeply alter their ability to mediate viral entry (36,37). A limitation of our study is that we were not always able to test the receptor orthologues of the actual bat species where the viruses were identified. The genomes of some bat species are not available and attempts of gene amplification from feces samples were often unsuccessful. Research efforts such as the Bat1K project (https://bat1k.com), which aims to sequence the genomes of all 1400 living bat species, will help to fill this knowledge gap and will allow to precisely assess the ability of a virus to use receptor orthologues of the bat species where it was identified.

Therefore, although we cannot completely exclude that the identified viruses may use ACE2, APN, DPP4 or TMPRSS2 from their host species as a receptor, it is likely that they use other yet unknown receptors for entry. Indeed, three of the identified viruses were minacoviruses and one was a nyctacovirus, and no receptor has been described yet for viruses of these subgenera. Finally, three of the identified alphacoronaviruses belonged to the pedacovirus subgenus, named after the porcine epidemic diarrhea virus (PEDV). Although PEDV has been suggested to use APN as a receptor (41,42), this has been debated recently as no specific interaction between the PEDV spike and APN could be measured (43,44) and APN knockout pigs are susceptible to PEDV infection (45). Therefore, PEDV and other bat-borne pedacoviruses may use a receptor other than APN.

In conclusion, our results highlight the role of bats as a global reservoir of a wide range of coronaviruses, including Spain. Future work aimed at characterizing the diversity of coronaviruses in different host types and regions, as well as ecological monitoring of interactions between bats and other species, including humans, will be crucial to mitigate the risks of future pandemics.

## Materials and methods

### Sample collection

To capture bats from their natural habitats, nylon mist nets (Ecotone) and harp traps (Austbat) were used in seven Spanish regions (Cantabria, Castellón, Lugo, Murcia, Salamanca, Teruel, and Valencia) between May and October 2022 (**Table S1, Figure 1**). Each captured animal was identified at the species level, sexed, measured, aged, and briefly placed in cotton bags to recover fresh fecal samples. Samples were obtained from 202 individuals representing 23 species of bats (19 from the *Vespertilionidae* family, 3 from the *Rhinolophidae* family and 1 from the *Molossidae* family). Exceptionally, fresh samples from European free-tailed bat (Tadarida teniotis) were collected from a colony without trapping involved. Fecal samples were placed in tubes containing 500 µL of phosphate-buffered saline (PBS) and maintained at 4ºC throughout the duration of the fieldwork (6 to 9 h). Samples were then kept at -20°C until arrival at the laboratory where they were stored at -80°C.

### Sample processing and RNA extraction

A subset of the fecal samples from each of the 202 individuals was combined into 26 pools, each one containing between 1 and 15 samples from the same bat species (**Table S1**). Prior to sample processing, each pool was spiked with 10^5^ plaque-forming units (PFU) of VSV as a positive control to assess the final viral recovery efficiency. Each tube was mixed with PBS, resulting in a final volume of 1.5 mL. Fecal samples were homogenized using the Precellys Evolution tissue homogenizer (Bertin) in 2 mL tubes with 1.4 mm ceramic beads (Precellys), using three cycles of 30 sec at 6500 rpm, with 10 sec pause between cycles. The homogenates were centrifuged twice at 20000 g for 3 min at 4°C. The resulting supernatants were filtered using a Minisart cellulose acetate syringe filter with a 1.2-µm pore size (Sartorius) and transferred to ultra-clean 2 mL tubes (Eppendorf). RNA extraction was performed using 280 µL of the total filtered volume with the QIAamp Viral RNA minikit (Qiagen). RNA was eluted in a final volume of 40 µL and stored at - 80°C.

### Sequencing and annotation

The extracted RNA was subjected to library preparation using the stranded mRNA preparation kit (Illumina) but starting at the fragmentation stage. Samples were subjected to paired-end sequencing using a NextSeq 550 device with read length of 150 bp at each end (**Table S2**). Raw reads were deduplicated, quality filtered with a quality trimming threshold of 20, and any reads below 70 nucleotides in length were removed using fastp v0.23.2 (46). *De novo* sequence assembly was performed using SPAdes v3.15.4 with the meta option (47), as well as using MEGAHIT v1.2.9 (48) with default parameters. Assembled contigs were clustered to remove replicates or small replicates of larger contigs, using CD-HIT v4.8.1 (49). Contigs shorter than 1000 nt were removed and the remaining sequences were taxonomically classified using Kaiju v1.9.0 (50) with the subset of the National Center for Biotechnology Information (NCBI) nr protein database comprising archaea, bacteria and viruses (downloaded on June 6, 2023). Then, all clustered sequences were analyzed using Virsorter2 v2.2.4 (51) to detect viral contigs. Following this, the quality of the viral contigs was further assessed using CheckV v1.0.1 with the CheckV database v1.5. Contigs corresponding to phages and those that could not be classified into a known viral family were excluded. The remaining contigs were selected based on their size, completeness, and the ability of the assigned virus family to infect vertebrates. Finally, contigs that were assigned to the *Coronaviridae* family were selected. Coverage statistics were obtained by remapping the trimmed and filtered reads to each contig using Bowtie2 v2.2.5 (52). Open reading frames (ORFs) were predicted using ORFfinder (ncbi.nlm.nih.gov/orffinder), while protein domains were annotated using InterProScan v5.63-95.0 (53) with the Pfam database v35.0.

### Accession numbers

The viral sequences presented in this study are accessible via the GenBank database, with the following accession numbers: PQ611137 (Sarbecovirus sp. isolate RhGB02 (variant Murcia2022)), PQ611138 (Myotis capaccinii associated Yeseras alphacoronavirus), PQ611139 (Myotis escalerai associated Sima alphacoronavirus), PQ611140 (Miniopterus schreibersii associated Gordo alphacoronavirus), PQ611141 (Alphacoronavirus sp. isolate ATG20-YN-CHN-2020 (variant Murcia2022)), PQ611142 (Alphacoronavirus Bat-CoV/P.kuhlii/Italy/206679-3/2010 (variant Valencia2022)). The novel bat gene sequences obtained in this study are also accessible via the GenBank database, with the following accession numbers: PQ611143 (*Miniopterus schreibersii* ACE2), PQ611154 (*Myotis escalerai* APN), PQ611150 (*Miniopterus schreibersii* APN allele 1), PQ611151 (*Miniopterus schreibersii* APN allele 2), PQ611152 (*Miniopterus schreibersii* TMPRSS2) and PQ611153 (*Myotis escalerai* TMPRSS2).

### Phylogenetic and taxonomical positioning

To place the sequences within the global coronavirus diversity, the *Coronaviridae* ICTV report (18) was used to identify representative species. RdRP and helicase amino acid sequences from representative coronavirus species were downloaded from the NCBI. The sequence of Microhyla letovirus 1 was included as an outgroup for comparative purposes. Sequences were aligned with Clustal Omega v1.2.3 (54) and a maximum likelihood (ML) tree was constructed under the LG+I+G4 substitution model. Phylogenetic analyses were conducted using IQTree v2.0.3 (55), and model selection was performed with the built-in ModelFinder feature (56). Branch support was estimated using ultra-fast boot-strapping replicates (UFBoot2) (57) and an approximate likelihood-ratio test (SH-aLRT) (58) with 1000 replicates. The resulting phylogenetic trees were visualized using iTol v6.0.9 (59). To assess whether the identified viruses represented novel species, the percentage of nucleotide sequence identity was obtained for the closest viral genomes using BLASTn (**Table S3**).

### RT-PCR amplification of viral sequences

RT-PCR was used to check for the presence of the identified coronaviruses in each of the individual samples within each positive pool (**Table S4**). RNA was extracted from each individual fecal sample using the Qiagen QIAamp Viral RNA minikit (Qiagen), following manufacturer’s instructions, and eluted in a final volume of 30 µL. Specific primers were designed for cDNA synthesis and subsequent PCR amplification of a 500 bp spike region of each virus of interest (**Table S5**). A volume of 4 µL of the RNA extraction was used for cDNA synthesis using Invitrogen’s Superscript IV enzyme (Invitrogen). For PCR, the NZYTaq II Green Master Mix (NZYTech) was used. Sample positivity was checked by electrophoresis on a 1% agarose gel using NZYTech Green Safe Premium (NZYTech).

### Phylogenetic analysis of spike protein sequences

To explore the evolutionary origin of the identified coronaviruses, we constructed spike phylogenetic trees of each subgenus using the closest amino acid sequences from each identified sequence selected with BLASTp (**Table S3**). Spike sequences were downloaded from the NCBI. The sequence of Microhyla letovirus 1 was included as an outgroup for comparative purposes. Sequences were aligned with Clustal Omega v1.2.3 (54) and a ML tree was constructed under the WAG+F+G4 substitution model. Phylogenetic analyses were conducted as previously described.

### Cell culture

HEK293T cells were obtained from the American Type Culture Collection (ATCC, CRL-3216) and cultured in Dulbecco’s Modified Eagle’s Medium (DMEM; Gibco) supplemented with 10% fetal bovine serum (FBS, Gibco), 1% non-essential amino acids (NEAA; Gibco), penicillin and streptomycin (P/S; 10 units/mL and 10 µg/mL, respectively; Gibco) and amphotericin B (250 ng/mL, Gibco). Huh7 cells were kindly provided by Francis Chisari and VeroE6 cells were obtained from the ATCC (ATCC-1586). Both were maintained in DMEM supplemented with 10% FBS (Sigma), 1% P/S (Gibco) and 1% NEAA (Gibco). All cells were maintained at 5% CO_2_ and 37°C in a humidified incubator and were routinely screened for the presence of mycoplasma by PCR.

### Viral isolation attempts

Fecal samples positive for the presence of the novel alphacoronaviruses (McAlphaCoV-Yeseras, MeAlphaCoV-Sima and MsAlphaCoV-Gordo) and the betacoronavirus (RhBetaCoV-Murcia2022) were used to inoculate VeroE6 and Huh-7 cells. Cells were seeded in 12-well plates to achieve an 80% confluence on the day of infection and incubated at 37°C with 5% CO_2_. The following day, cells were infected at 37°C for 2 h with two distinct conditions: (I) 100 µL of viral samples and (II) 100 µL of viral samples with 2 µg/mL trypsin (T1426; Sigma-Aldrich). Subsequently, 1 mL of DMEM supplemented with 10% FBS and antibiotics was added to each well, and cells were incubated at 37ºC with 5% CO_2_. After 96 h, cells and supernatants (cleared by centrifugation at 2000 g for 10 min) were collected, aliquoted, and stored at -80°C. Viral RNA from the supernatants was extracted using the *Quick*-RNA Viral Kit (Zymo Research), following manufacturer’s instructions. Viral RNA from cells was extracted using phenol-chloroform. Extracted RNA was initially reverse transcribed (RT) using the SuperScript III Reverse Transcriptase (Invitrogen) and the resulting cDNA was used in a qPCR using the SYBR Green PowerUp (Applied Biosystems) and specific primers (**Table S6**). Supernatants of the McAlphaCoV-Yeseras and RhBetaCoV-Murcia2022 samples collected from the initial infection were used to initiate a second passage. Supernatants from the two infection conditions (with and without trypsin treatment) were pooled and used for infecting the same cell line in which they were collected. After 3 and 6 days, cells and supernatants were collected, centrifuged at 2000g for 10 min, aliquoted, and stored at -80ºC. Viral RNA was extracted from cells and supernatants and processed as previously described. Probe-based RT-qPCR was conducted using the TaqPath 1-Step RT-qPCR Master Mix (ThermoFisher) and specific primers/probes (**Table S6**).

### Viral pseudotyping

Human codon-optimized spike constructs of the six complete coronaviruses identified were ordered as synthetic genes and cloned in a pcDNA3.1-C-HisTag vector (Genscript). For pseudotyping, T75 flasks were coated with poly-D-lysine (Gibco) for 2 h at 37°C, washed with distilled water, and seeded with 8 × 10^6^ HEK293T cells. The following day, cells were transfected with 30 µg of viral glycoprotein expression plasmid using Lipofectamine 2000 (Invitrogen) following manufacturer’s instructions. To produce negative control bald pseudotypes, cells were transfected with an empty pcDNA3.1 vector. At 24 h post-transfection, cells were inoculated at a multiplicity of infection (MOI) of 3 infectious units per cell for 1 h at 37°C with a VSV encoding GFP, lacking the glycoprotein gene G (VSVΔG-GFP), and previously pseudotyped with G. Following this, cells were washed three times with PBS and 8 mL of DMEM supplemented with 2% FBS were added. Supernatants were harvested 24 h later, cleared by centrifugation at 2000 g for 10 min, passed through a 0.45 µm filter, aliquoted, and stored at -80°C.

### Cloning of human and bat genes

The human ACE2 (hACE2)-encoding plasmid was kindly provided by Dr. Ron Geller (I2SysBio-CSIC). For each gene of interest, the sequence of the main transcript was retrieved from the NCBI RefSeq (ncbi.nlm.nih.gov/refseq), UniProt (uniprot.org/uniprotkb) or Ensembl (ensembl.org) databases, when available. The ACE2 CDS sequence from *Rhinolophus sinicus* (AGZ48803.1), *Rhinolophus cornutus* (BCG67443.1), *Myotis myotis* (XM_036305841.1) and *Pipistrellus kuhlii* (XM_036439529.2) and the APN gene sequence from *Pipistrellus kuhlii* (XM_036415540.2) were ordered as synthetic genes in a pcDNA3.1-C-FlagTag vector (GenScript). The human CDS sequences of APN (hAPN, NM_001150.3), DPP4 (hDPP4, NM_001935.4) and TMPRSS2 (hTMPRSS2, NM_005656.4) were obtained from RNA extracted using RNAzol (Sigma-Aldrich) from the CAKI-1, TK-10 and SW-620 cell lines, respectively. The ACE2 sequence (ACE2) from *Miniopterus schreibersii*, the APN sequence from *Myotis escalerai* and *Miniopterus schreibersii*, and the TMPRSS2 sequence from *Miniopterus schreibersii* and *Myotis escalerai* bat were obtained from the RNA extracted from fecal samples of the respective bat species using the NZY Total RNA Isolation kit (NZYtech). RNA was reverse transcribed into cDNA using the SuperScript IV Reverse Transcriptase (Invitrogen) and specific primers (**Table S7**) following the manufacturer’s instructions. CDSs were cloned into a pcDNA3.1-C-FlagTag vector via HiFi assembly. Briefly, the pcDNA3.1-C-FlagTag vector was linearized by PCR (Forward primer: 5’-GATTACAAGGATGACGACGATAAGTG-3’; Reverse primer: 5’-GGTGGCAAGCTTAAGTTTAAACGCTAG-3’) and CDSs were amplified from cDNAs using specific primers (**Table S7**) that contained a 20-nucleotide tail overlapping with the 5’ or 3’ ends of the linearized pcDNA3.1-C-Flag vector. The linearized vector and PCR-amplified sequences were purified using the DNA Clean & Concentrator-5 kit (Zymo Research), mixed in a 1:2 molar ratio, and assembled using the NEBuilder HiFi DNA Assembly Master Mix (New England Biolabs), following manufacturer’s instructions. All PCR steps were conducted using Phusion Hot Start II High-Fidelity DNA polymerase (Thermo Scientific). Assembled products were transformed into NY5α competent cells (NZYtech). Correct insertion was verified through colony PCR, using vector-specific primers (Forward primer: 5’-GAGAACCCACTGCTTACTGGC-3’; Reverse primer: 5’-AGGGTCAAGGAAGGCACG-3’) and the NZYTaq II 2x Green Master Mix (NZYtech). Plasmids were checked by whole-plasmid high-throughput sequencing (Plasmidsaurus or Eurofins). Protein expression was confirmed by western blot or pseudotype infection (see below).

### Western blot

Viral pseudotypes were pelleted by centrifugation at 30000 g for 2 h at 4°C. The pellet was lysed in 30 µL of NP-40 lysis buffer (Invitrogen) supplemented with a complete protease inhibitor (Roche) and incubated for 30 min on ice. Transfected cells were lysed in 100 µL of NP-40 lysis buffer supplemented with a complete protease inhibitor (Roche) and incubated for 30 min on ice. The lysates were then cleared by centrifugation at 15000 g for 10 min at 4°C. Viral and cellular lysates were mixed with 4x Laemmli buffer (Bio-Rad) supplemented with 10% β-mercaptoethanol and denatured at 95°C for 5 min. Proteins were separated by SDS-PAGE using pre-cast 4-20% Mini-PROTEAN TGX Gels (Bio-Rad) and transferred onto a 0.45 µm PVDF membrane (Thermo Scientific). Membranes were blocked with TBS-T (20 mM tris, 150 nM NaCl, 0.1% Tween-20, pH 7.5) supplemented with 3% bovine serum albumin (BSA; Sigma) for 1 h at room temperature (RT). Membranes were then incubated for 1 h at room temperature with the following primary antibodies: rabbit anti-His-Tag (dilution 1:1000, Invitrogen PA1-983B), mouse anti-VSV-M (dilution 1:1000, clone 23H12, Kerafast EB0011), mouse anti-Flag-Tag (dilution 1:1000, clone M2, Sigma-Aldrich F1804) and rabbit anti-GAPDH (dilution 1:3000, Sigma-Aldrich ABS16). After three washes with TBS-T, the primary antibody was detected using a horseradish peroxidase (HRP)-conjugated anti-mouse (dilution 1:50000, Invitrogen, G-21040) or anti-rabbit (dilution 1:50000, Invitrogen, G-21234) secondary antibody. After three washes in TBS-T, signal was revealed using SuperSignal™ West Pico PLUS (Thermo Scientific), following manufacturer’s instructions. Images were captured using an ImageQuant LAS 500 (GE Healthcare) and analyzed using Fiji software v.2.14.0.

### Pseudotype infection assays

HEK293T cells were seeded in 96-well plates (3.5 × 10^4^ cells per well) and incubated at 37°C, 5% CO_2_ for 24 h. The following day, cells were transfected with 100 ng of plasmids encoding the indicated human and bat orthologues of ACE2, DPP4, APN and TMPRSS2, or an empty vector, using Lipofectamine 2000 (Invitrogen). After 24 h, spike-expressing pseudotypes were mixed 1:1 with a house-made anti-VSV-G monoclonal antibody and incubated for 20 min at 37°C. Cell culture medium was then removed, and cells were inoculated with 50 µL of the antibody-treated pseudotypes. After 2 h at 37°C, 50 µL of DMEM supplemented with 2% FBS was added to each well. After 24 h, plates were imaged in an Incucyte SX5 Live-Cell Analysis System (Sartorius). Cell confluence and the percentage of GFP-positive area were quantified automatically with the Incucyte Analysis software to determine the percentage of infected cells.

## Acknowledgements

We thank all members of the Virus Evolution laboratory for helpful discussions about this work. We thank Xose Pardavila (Sorex), Jorge Sereno-Cadierno, Morcegos de Galicia, Raul Molleda, Sandra Córdoba, and Ana Cordero for their collaboration during the bat-trapping and sample collection. This research was financially supported by an ERC Advanced Grant (101019724— EVADER), grant PID2020-118602RB-I00 from the Spanish Ministerio de Ciencia e Innovación (MICINN), and grant CIAICO/2022/110 from the Conselleria de Educación, Universidades y Empleo (Generalitat Valenciana) to R.S. J.Du. was supported by an EMBO postdoctoral fellowship (ALTF 140-2021) and a Marie Skłodowska-Curie Actions Postdoctoral Fellowship (101104880).

## Ethics statement

Samples consisted of feces from wild animals captured using nylon mist nets or a harp trap. Bats were kept briefly in cotton bags until fresh fecal samples were obtained. According to the European Directive regulating the protection of animals used for scientific purposes (2010/62/EU, Article 1), subsequently transposed into Spanish legislation (Royal Decree 53/213, 1 February, Article 2), procedures used in this study are not subject to the condition of animal experimentation and therefore do not require approval by an institutional ethics committee, but specifically a permit for fieldwork from the competent regional authority. The necessary permits from the Generalitat Valenciana for the sampling of wild bats were granted under Exp. 2022-VS (FAU22_009).

## Author contributions

J.M.C., J.Du. and R.S. designed research; A.V. collected samples; C.S.-T., E.M., R.M.-R. performed research; J.B., C.S.-T., E.M., J.Du. and R.S. analyzed data; J.S.M., J. Dí, J.M.C., J.Du., R.S. supervised work; C.S.-T., J.Du. and R.S. wrote the manuscript; J.M.C. and R.S. provided funding. All authors read and approved the final version of the manuscript.

## References

1. Letko M, Seifert SN, Olival KJ, Plowright RK, Munster VJ. Bat-borne virus diversity, spillover and emergence. Vol. 18, Nature Reviews Microbiology. Nature Research; 2020. p. 461–71.

2. Calisher CH, Childs JE, Field HE, Holmes K V., Schountz T. Bats: Important reservoir hosts of emerging viruses. Vol. 19, Clinical Microbiology Reviews. 2006. p. 531–45.

3. Drexler JF, Corman VM, Gloza-Rausch F, Seebens A, Annan A, Ipsen A, et al. Henipavirus RNA in African bats. PLoS One. 2009 Jul 28;4(7).

4. Leroy E, Kumulungui B, Pourrut X, et al. Fruit bats as reservoirs of Ebola virus. Nature. 2005;438:575–6.

5. Li W, Shi Z, Yu M, Ren W, Smith C, Epstein JH, et al. Bats Are Natural Reservoirs of SARS-Like Coronaviruses. Science [Internet]. 2005;310(5748):676–9. Available from: http://www.cdc.gov/ncidod/EID/vol11no10/05-0513.htm

6. Anthony SJ, Gilardi K, Menachery VD, Goldstein T, Ssebide B, Mbabazi R, et al. Further evidence for bats as the evolutionary source of middle east respiratory syndrome coronavirus. mBio. 2017 Mar 1;8(2).

7. Hu B, Ge X, Wang LF, Shi Z. Bat origin of human coronaviruses Coronaviruses: Emerging and re-emerging pathogens in humans and animals Susanna Lau Positive-strand RNA viruses. Vol. 12, Virology Journal. BioMed Central Ltd.; 2015.

8. Mollentze N, Streicker DG. Viral zoonotic risk is homogenous among taxonomic orders of mammalian and avian reservoir hosts. PNAS. 2020;117(17):9423–30.

9. Wong S, Lau S, Woo P, Yuen KY. Bats as a continuing source of emerging infections in humans. Rev Med Virol. 2007 Mar;17(2):67–91.

10. Waruhiu C, Ommeh S, Obanda V, Agwanda B, Gakuya F, Ge XY, et al. Molecular detection of viruses in Kenyan bats and discovery of novel astroviruses, caliciviruses and rotaviruses. Virol Sin. 2017 Apr 1;32(2):101–14.

11. Rizzo F, Edenborough KM, Toffoli R, Culasso P, Zoppi S, Dondo A, et al. Coronavirus and paramyxovirus in bats from Northwest Italy. BMC Vet Res. 2017 Dec 22;13(1).

12. Crook JM, Murphy I, Carter DP, Pullan ST, Carroll M, Vipond R, et al. Metagenomic identification of a new sarbecovirus from horseshoe bats in Europe. Sci Rep. 2021 Dec 1;11(1).

13. Tan CCS, Trew J, Peacock TP, Mok KY, Hart C, Lau K, et al. Genomic screening of 16 UK native bat species through conservationist networks uncovers coronaviruses with zoonotic potential. Nat Commun. 2023 Dec 1;14(1).

14. Zhang YZ, Shi M, Holmes EC. Using Metagenomics to Characterize an Expanding Virosphere. Vol. 172, Cell. Cell Press; 2018. p. 1168–72.

15. Bassi C, Guerriero P, Pierantoni M, Callegari E, Sabbioni S. Novel Virus Identification through Metagenomics: A Systematic Review. Vol. 12, Life. MDPI; 2022.

16. Ruiz-Aravena M, McKee C, Gamble A, Lunn T, Morris A, Snedden CE, et al. Ecology, evolution and spillover of coronaviruses from bats. Nat Rev Microbiol. 2022;20(5):299– 314.

17. Cui J, Li F, Shi ZL. Origin and evolution of pathogenic coronaviruses. Vol. 17, Nature Reviews Microbiology. Nature Publishing Group;2019. p. 181–92.

18. Woo PCY, De Groot RJ, Haagmans B, Lau SKP, Neuman BW, Perlman S, et al. ICTV Virus Taxonomy Profile: Coronaviridae 2023. Journal of General Virology. 2023;104(4).

19. Graham RL, Baric RS. Recombination, Reservoirs, and the Modular Spike: Mechanisms of Coronavirus Cross-Species Transmission. J Virol. 2010 Apr;84(7):3134–46.

20. Temmam S, Vongphayloth K, Baquero E, Munier S, Bonomi M, Regnault B, et al. Bat coronaviruses related to SARS-CoV-2 and infectious for human cells. Nature. 2022 Apr 14;604(7905):330–6.

21. Dufloo J, Andreu-Moreno I, Valero-Rello A, Sanjuán R. Viral entry is a weak barrier to zoonosis. bioRxiv [Internet]. 2024 Jan 23 [cited 2024 May 29];2024.01.22.576693. Available from: https://www.biorxiv.org/content/10.1101/2024.01.22.576693v1

22. Chen L, Liu B, Yang J, Jin Q. DBatVir: The database of bat-associated viruses. Database. 2014;2014.

23. Falcón A, Vázquez-Morón S, Casas I, Aznar C, Ruiz G, Pozo F, et al. Detection of alpha and betacoronaviruses in multiple Iberian bat species. Arch Virol. 2011;156(10):1883– 90.

24. Moraga-Fernández A, Sánchez-Sánchez M, Queirós J, Lopes AM, Vicente J, Pardavila X, et al. A study of viral pathogens in bat species in the Iberian Peninsula: identification of new coronavirus genetic variants. Int J Vet Sci Med. 2022;10(1):100–10.

25. Buigues J, Viñals A, Martínez-Recio R, Monrós JS, Cuevas JM, Sanjuán R. Phylogenetic evidence supporting the nonenveloped nature of hepadnavirus ancestors. Proc Natl Acad Sci U S A. 2024;121(45).

26. Buigues J, Viñals A, Martínez-Recio R, Monrós JS, Sanjuán R, Cuevas JM. Full-genome sequencing of dozens of new DNA viruses found in Spanish bat feces. He B, editor. Microbiol Spectr. 2024;0(0).

27. Carrascosa-Sàez M, Buigues J, Viñals A, Andreu-Moreno I, Martínez-Recio R, Soriano-Tordera C, et al. Genetic diversity and cross-species transmissibility of bat-associated picornaviruses from Spain. Virol J. 2024;21(1):193.

28. Nayfach S, Camargo AP, Schulz F, Eloe-Fadrosh E, Roux S, Kyrpides NC. CheckV assesses the quality and completeness of metagenome-assembled viral genomes. Nat Biotechnol. 2021 May 1;39(5):578–85.

29. Alkhovsky S, Lenshin S, Romashin A, Vishnevskaya T, Vyshemirsky O, Bulycheva Y, et al. SARS-Like Coronaviruses in Horseshoe Bats (Rhinolophus spp.) in Russia, 2020. Viruses. 2022 Jan 1;14(1).

30. Vijaykrishna D, Smith GJD, Zhang JX, Peiris JSM, Chen H, Guan Y. Evolutionary Insights into the Ecology of Coronaviruses. J Virol. 2007 Apr 15;81(8):4012–20.

31. Han Z, Xiao J, Song Y, Zhao X, Sun Q, Lu H, et al. Highly diverse ribonucleic acid viruses in the viromes of eukaryotic host species in Yunnan province, China. Front Microbiol. 2022 Oct 13;13.

32. De Sabato L, Lelli D, Faccin F, Canziani S, Di Bartolo I, Vaccari G, et al. Full genome characterization of two novel Alpha-coronavirus species from Italian bats. Virus Res. 2019 Jan 15;260:60–6.

33. Han Y, Xu P, Wang Y, Zhao W, Zhang J, Zhang S, et al. Panoramic analysis of coronaviruses carried by representative bat species in Southern China to better understand the coronavirus sphere. Nat Commun. 2023;14:5537(1).

34. Strizhakova O, Hanke D, Titov I, Blome S, Malogolovkin A. Complete genome sequence of a porcine epidemic diarrhea virus isolated in Belgorod, Russia, in 2008. Genome Announc. 2017;5(41).

35. Wiederkehr MA, Qi W, Schoenbaechler K, Fraefel C, Kubacki J. Virus Diversity, Abundance, and Evolution in Three Different Bat Colonies in Switzerland. Viruses. 2022 Sep 1;14(9).

36. Roelle SM, Shukla N, Pham AT, Bruchez AM, Matreyek KA. Expanded ACE2 dependencies of diverse SARS-like coronavirus receptor binding domains. PLoS Biol. 2022 Jul 1;20(7).

37. Starr TN, Zepeda SK, Walls AC, Greaney AJ, Alkhovsky S, Veesler D, et al. ACE2 binding is an ancestral and evolvable trait of sarbecoviruses. Nature. 2022 Mar 31;603(7903):913–8.

38. Si JY, Chen YM, Sun YH, Gu MX, Huang ML, Shi LL, et al. Sarbecovirus RBD indels and specific residues dictating multi-species ACE2 adaptiveness. Nat Commun [Internet]. 2024 Oct 14;15(1):8869. Available from: https://www.nature.com/articles/s41467-024-53029-3

39. Caraballo DA. Cross-Species Transmission of Bat Coronaviruses in the Americas: Contrasting Patterns between Alphacoronavirus and Betacoronavirus. Microbiol Spectr. 2022 Aug 31;10(4).

40. Latinne A, Hu B, Olival KJ, Zhu G, Zhang L, Li H, et al. Origin and cross-species transmission of bat coronaviruses in China. Nat Commun. 2020 Dec 1;11(1).

41. Li BX, Ge JW, Li YJ. Porcine aminopeptidase N is a functional receptor for the PEDV coronavirus. Virology. 2007 Aug 15;365(1):166–72.

42. Liu C, Tang J, Ma Y, Liang X, Yang Y, Peng G, et al. Receptor Usage and Cell Entry of Porcine Epidemic Diarrhea Coronavirus. J Virol. 2015 Jun;89(11):6121–5.

43. Li W, Luo R, He Q, van Kuppeveld FJM, Rottier PJM, Bosch BJ. Aminopeptidase N is not required for porcine epidemic diarrhea virus cell entry. Virus Res. 2017 May 2;235:6– 13.

44. Shirato K, Maejima M, Islam MT, Miyazaki A, Kawase M, Matsuyama S, et al. Porcine aminopeptidase N is not a cellular receptor of porcine epidemic diarrhea virus, but promotes its infectivity via aminopeptidase activity. Journal of General Virology. 2016 Oct 1;97(10):2528–39.

45. Zhang J, Wu Z, Yang H. Aminopeptidase N Knockout Pigs Are Not Resistant to Porcine Epidemic Diarrhea Virus Infection. Vol. 34, Virologica Sinica. Science Press; 2019. p. 592–5.

46. Chen S, Zhou Y, Chen Y, Gu J. Fastp: An ultra-fast all-in-one FASTQ preprocessor. In: Bioinformatics. Oxford University Press; 2018. p. i884–90.

47. Nurk S, Meleshko D, Korobeynikov A, Pevzner PA. MetaSPAdes: A new versatile metagenomic assembler. Genome Res. 2017 May 1;27(5):824–34.

48. Li D, Liu CM, Luo R, Sadakane K, Lam TW. MEGAHIT: An ultra-fast single-node solution for large and complex metagenomics assembly via succinct de Bruijn graph. Bioinformatics. 2015 May 15;31(10):1674–6.

49. Li W, Godzik A. Cd-hit: A fast program for clustering and comparing large sets of protein or nucleotide sequences. Bioinformatics. 2006 Jul 1;22(13):1658–9.

50. Menzel P, Ng KL, Krogh A. Fast and sensitive taxonomic classification for metagenomics with Kaiju. Nat Commun. 2016 Apr 13;7.

51. Guo J, Bolduc B, Zayed AA, Varsani A, Dominguez-Huerta G, Delmont TO, et al. VirSorter2: a multi-classifier, expert-guided approach to detect diverse DNA and RNA viruses. Microbiome. 2021 Dec 1;9(1).

52. Langmead B, Salzberg SL. Fast gapped-read alignment with Bowtie 2. Nat Methods. 2012 Apr;9(4):357–9.

53. Jones P, Binns D, Chang HY, Fraser M, Li W, McAnulla C, et al. InterProScan 5: Genome-scale protein function classification. Bioinformatics. 2014 May 1;30(9):1236–40.

54. Sievers F, Wilm A, Dineen D, Gibson TJ, Karplus K, Li W, et al. Fast, scalable generation of high-quality protein multiple sequence alignments using Clustal Omega. Mol Syst Biol. 2011;7.

55. Minh BQ, Schmidt HA, Chernomor O, Schrempf D, Woodhams MD, Von Haeseler A, et al. IQ-TREE 2: New Models and Efficient Methods for Phylogenetic Inference in the Genomic Era. Mol Biol Evol. 2020 May 1;37(5):1530–4.

56. Kalyaanamoorthy S, Minh BQ, Wong TKF, Von Haeseler A, Jermiin LS. ModelFinder: Fast model selection for accurate phylogenetic estimates. Nat Methods. 2017 May 30;14(6):587–9.

57. Thi Hoang D, Chernomor O, von Haeseler A, Quang Minh B, Sy Vinh L, Rosenberg MS. UFBoot2: Improving the Ultrafast Bootstrap Approximation. Mol Biol Evol. 2017;35(2):518–22.

58. Guindon S, Dufayard JF, Lefort V, Anisimova M, Hordijk W, Gascuel O. New algorithms and methods to estimate maximum-likelihood phylogenies: Assessing the performance of PhyML 3.0. Syst Biol. 2010 May;59(3):307–21.

59. Letunic I, Bork P. Interactive Tree of Life (iTOL) v6: Recent updates to the phylogenetic tree display and annotation tool. Nucleic Acids Res. 2024 Jul 5;52(W1):W78–82.

